# Quantifying the strength of viral fitness tradeoffs between hosts: A meta-analysis of pleiotropic fitness effects

**DOI:** 10.1101/2023.12.16.571995

**Authors:** Xuechun ‘May’ Wang, Julia Muller, Mya McDowell, David A. Rasmussen

## Abstract

The range of hosts a given virus can infect is widely presumed to be limited by fitness tradeoffs between alternative hosts. These fitness tradeoffs may arise naturally due to antagonistic pleiotropy if mutations that increase fitness in one host tend to decrease fitness in alternate hosts. Yet there is also growing recognition that positive pleiotropy may be more common than previously appreciated. With positive pleiotropy, mutations have concordant fitness effects such that a beneficial mutation can simultaneously increase fitness in different hosts, providing a genetic mechanism by which selection can overcome fitness tradeoffs. How readily evolution can overcome fitness tradeoffs therefore depends on the overall distribution of mutational fitness effects between hosts, including the relative frequency of antagonistic versus positive pleiotropy. We therefore conducted a systematic meta-analysis of the pleiotropic fitness effects of viral mutations reported in different hosts. Our analysis indicates that while both antagonistic and positive pleiotropy are common, fitness effects are overall positively correlated between hosts and unconditionally beneficial mutations are not uncommon. Moreover, the relative frequency of antagonistic versus positive pleiotropy may simply reflect the underlying frequency of beneficial and deleterious mutations in individual hosts. Given a mutation is beneficial in one host, the probability that it is deleterious in another host is roughly equal to the probability that any mutation is deleterious, suggesting there is no natural tendency towards antagonistic pleiotropy. The widespread prevalence of positive pleiotropy suggests that many fitness tradeoffs may be readily overcome by evolution given the right selection pressures.

Lay summary

Evolutionary theory suggests that fitness tradeoffs between alternative environments constrain the potential for organisms to simultaneously adapt to multiple environments. Likewise, fitness tradeoffs between alternative hosts are widely believed to limit the ability of viruses to adapt to multiple hosts and thereby expand their host range. How strongly viruses are constrained by such tradeoffs will largely depend on the fitness effects of new mutations. Fitness tradeoffs may inevitably constrain viral evolution if mutations that increase fitness in one host tend to decrease fitness in alternative hosts. However, mutations can sometimes increase fitness in multiple hosts, allowing viruses to adapt to new hosts without paying fitness costs. Geneticists refer to these two scenarios as antagonistic and positive pleiotropy depending on whether mutations have opposite or concordant fitness effects. Because the relative frequency of antagonistic versus positive pleiotropy is centrally important to viral evolution, we conducted a systematic meta-analysis of the fitness effects of mutations reported in different hosts. Our analysis reveals that cases of positive pleiotropy where mutations have beneficial effects in more than one host may be sufficiently common for evolution to resolve many apparent fitness tradeoffs between hosts.

## Introduction

Viruses regularly jump between hosts and exhibit a tremendous capacity to adapt to novel host environments (Geoghegan et al., 2017; Longdon et al., 2014). Consequently, new viral pathogens regularly emerge on novel hosts and expand their host range (Lloyd-Smith et al., 2009). Nevertheless, many observations suggest that the host range of individual viruses is restricted, often to a set of closely related hosts (Bedhomme et al., 2015; Dawson & Hilf, 1992; Longdon et al., 2011). Likewise, when viruses are experimentally passaged on a single host, fitness tends to decline rapidly in alternative hosts not encountered during passaging (Bedhomme et al., 2012; Kassen, 2002; Visher et al., 2022; Weaver et al., 1999). This seemingly paradoxical pattern of tremendous adaptive potential yet relatively restricted host ranges suggests that fitness tradeoffs may constrain the ability of viruses to adapt to multiple hosts simultaneously, thereby limiting the emergence of generalist viruses.

Evolutionary theory emphasizes the role of antagonistic pleiotropy in generating fitness tradeoffs between hosts (Bedhomme et al., 2015; Duffy et al., 2000; Elena et al., 2009). Broadly defined, pleiotropy refers to an individual gene or mutation affecting multiple phenotypic traits or the same trait in different environments (Remold, 2012; Wagner & Zhang, 2011). Antagonistic pleiotropy results when a given mutation has opposite effects on two traits, such as a viral mutation being beneficial in one host but deleterious in another. By contrast, positive pleiotropy results when a mutation has concordant effects across traits or environments. In viruses, there is substantial evidence for individual mutations having antagonistic or opposite fitness effects between hosts (Coffey et al., 2008; Duffy et al., 2000; Lalić et al., 2011). For example, mutations in viral glycoproteins frequently exhibit antagonistic pleiotropy due to the need for these proteins to bind to highly host-specific receptors during cell entry (Greene et al., 2005; Rogers et al., 1983; Tsetsarkin et al., 2007; Urbanowicz et al., 2016). Antagonistic pleiotropy therefore provides a simple genetic explanation for fitness tradeoffs: if most mutations have antagonistic effects then gaining the ability to infect a new host will come at the cost of becoming less well adapted to alternate hosts.

If all mutations have antagonistic effects on fitness, then fitness tradeoffs would be absolute or “hard” in the sense that there would be no way for evolution to resolve tradeoffs since increasing fitness in one host would inevitably decrease fitness in an alternate host. Yet fitness tradeoffs are not always absolute and viruses can evolve to overcome tradeoffs. For example, generalists with high fitness across multiple hosts often evolve when viruses are alternately transferred between different hosts, even though fitness on alternate hosts tends to decline when viruses are passaged exclusively on a single host (Bedhomme et al., 2012; Coffey & Vignuzzi, 2010; Kassen, 2002; Smith-Tsurkan et al., 2010; Weaver et al., 1999). Moreover, viruses adapted to more than one host sometimes have increased fitness relative to specialists adapted to a single host, leading to what has been referred to as fitness trade-ups or no-cost generalists (see Burmeister & Turner, 2020; McGee et al., 2014). These fitness trade-ups can arise due to unconditionally beneficial mutations increasing fitness in more than one host simultaneously, a form of positive pleiotropy which may be more common than previously appreciated (Lalić et al., 2011; Moreno-Pérez et al., 2016; Ruark-Seward et al., 2020; Vale et al., 2012). Taken together, these observations suggest that many fitness tradeoffs may be “soft” in the sense that evolution could resolve the tradeoff given appropriate genetic variation and selection pressures. The strength of a fitness tradeoff in terms of how easy or difficult it will be for evolution to overcome will therefore largely depend on the relative frequency of antagonistic versus positive pleiotropy.

Given the central importance of pleiotropy to fitness tradeoffs and host range evolution, surprisingly little work has been done to quantify the overall distribution of pleiotropic fitness effects between hosts. We therefore conducted a systematic meta-analysis of published studies in which the fitness effects of individual viral mutations were estimated in two or more hosts, allowing us to quantify both the magnitude and sign of pleiotropy, including the frequency of antagonistic versus positive pleiotropy. We also included several viral and host features in our meta-analysis, allowing us to explore whether any features can predict the pleiotropic effects of mutations between different hosts. We find that fitness effects are overall positively correlated between hosts with positive pleiotropy being slightly more frequent than antagonistic pleiotropy. Perhaps more surprisingly, antagonistic pleiotropy is no more common than would be expected by chance given the frequency of beneficial versus deleterious mutations in individual hosts. We end by discussing the implications of these findings on the strength of fitness tradeoffs between hosts more generally.

## Methods

### Literature search

We searched for published studies reporting mutational fitness effects in viruses using Google Scholar and Web of Science with the search terms “virus* OR viral AND mutation* AND fitness”. However, the vast majority of studies returned only measured viral fitness in one host. We therefore added at least one of the following terms to our original search term: “pleiotropy OR pleiotropic”, “host range”, “host alternation”, “tradeoff”. Systematic searches were performed originally in January 2021 and then again in April 2023. Study relevance was initially determined based on title and abstract, which resulted in 180 relevant publications. Relevant studies cited by these initially identified papers were subsequently added. After examining the full text of each study, 26 remained that provided relevant data and met all inclusion criteria. Supplemental Table S1 provides a complete list of all included studies. From each study, we extracted the fitness effects of each viral mutation that was measured in two or more hosts. Because many studies report the fitness effects of more than one mutation or fitness effects in more than a single pair of hosts, multiple pairwise comparisons of fitness are often available from a single study. Wherever possible, we adhered to the PRISMA guidelines for meta-analysis in ecology and evolutionary biology (O’Dea et al., 2021). How our study addressed each of these guidelines is outlined in Supplemental Table S2.

### Study inclusion criteria

We only considered published studies which reported the fitness effects of individual mutations in DNA and RNA viruses. To be included, the fitness effects of a viral mutation had to be reported in at least two host environments. Different genotypes and ecotypes of the same host species were considered to be different hosts. Fitness effects determined both *in vivo* and using *in vitro* cell/tissue culture experiments were included, but we did not include fitness measured in extracellular environments. We included fitness measurements from competition assays as well as experiments that determined the fitness of each genotype in isolation. We only considered fitness effects determined in isogenic (single-mutant) backgrounds and did not include the fitness of any genotype with more than a single mutation relative to the wildtype. Because it is often unclear whether mutations were naturally occurring or experimentally derived, we included both and do not differentiate between the two. We only included studies that reported fitness in terms of viral replication or growth rates and excluded studies that used less direct proxies of fitness such as host virulence or disease symptoms.

### Quantifying and standardizing fitness effects

In order to quantify the fitness effect of a mutation in a given host, the fitness of the mutant genotype (*w_mut_*) was compared against the fitness of a wildtype/reference genotype (*w_ref_*).. If fitness effects were not reported as numerical values, we extracted data from figures using WebPlotDigitizer 4.4 (Rohatgi, n.d.) or, in one case, from gel band intensities using ImageJ (Rasband and contributors, National Institutes of Health, USA).

In order to meaningfully compare mutational fitness effects between hosts and across studies, we converted all reported values to a relative fitness ratio:

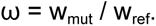

We chose to standardize fitness effects as ratios rather than differences because fitness differences depend on comparing rates in units of time that are often not transparent, complicating direct comparisons of fitness (Marée et al., 2000; Sanjuán, 2010; Sanjuán et al., 2004). Since that magnitude of fitness effects varied considerably between studies, we further log_2_ transform ω.

However, many studies report fitness in terms of selection coefficients (*s*) instead of the growth rates *w_mut_* and *w_ref_*. In this case, there is a straightforward relationship with growth rates: *s = w_mut_ - w_ref_* (Chevin, 2011; Fisher, 1930). We therefore simply convert selection coefficients to fitness ratios using the reported fitness of either genotype, e.g. ω = (s + w_ref_) / w_ref_. Alternatively, selection coefficients can be defined in terms of growth rates per discrete generation:

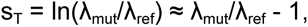

where *λ_mut_* and *λ_ref_* are the per-generation, multiplicative growth rates of the mutant and reference respectively. In this case, *s_T_* can be converted back to a fitness ratio using the relation ω *= s_T_ + 1* (Chevin, 2011; Sanjuán, 2010).

### Converting viral titers/abundance to fitness

Often no direct measure of fitness is reported but can be computed from growth curves showing changes in viral titers or abundance through time. In this case, we computed growth rates *w* from the change in viral abundance through time:

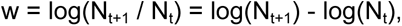

where *N_t_* and *N_t+1_* are the viral titers at two consecutive time points. If viral titres were reported at multiple time points, we computed the growth rate over each time interval and then recorded the average growth rate. In the later case, we only considered time points in the exponential growth phase up to the point at which viral titers peaked.

### Explanatory features

We recorded metadata about each mutation including features that might explain observed fitness effects. This included the mutation substitution type (synonymous vs. nonsynonymous), virus genome type (ssDNA, (+)ssRNA, (-)ssRNA, dsRNA), viral family, viral protein functional class (non-structural, structural, accessory), viral envelope status (present/absent), host type (plant, animal, bacteria), viral genome size and whether or not viral protein is involved in host-cell binding. Hereafter we refer to all these as explanatory features for simplicity. Additionally, we recorded the median estimated divergence time between each unique pair of hosts reported on timetree.org (Kumar et al., 2022).

### Quantifying pleiotropy

We differentiate between the magnitude and the sign of pleiotropy, following the terminology used to classify different types of epistasis (S. Remold, 2012; Weinreich et al., 2005). The sign of pleiotropy refers to whether a mutation has concordant (positive pleiotropy) or opposite (antagonistic pleiotropy) fitness effects between pairs of hosts. Note that our definition of positive pleiotropy encompasses all mutations with concordant sign effects, including unconditionally beneficial and unconditionally deleterious mutations. The magnitude of pleiotropy refers to the absolute difference in fitness effects between two hosts: *m = |ω_Host1_ - ω_Host2_|*. The sign and the magnitude of pleiotropy were computed for every unique pair of hosts in which fitness effects for a given mutation were available. Within each host pair, the ordering of hosts as *Host_1_* or *Host_2_*was randomly assigned as there is generally no way to determine which host is ancestral.

### Statistical analysis

The relationship between mutational fitness effects and explanatory features were explored using different linear regression models in *statsmodels* version 0.14 (Seabold & Perktold, 2010). We used a mixed effect linear model to test for significant associations between explanatory features (fixed effects) and fitness effects in single hosts. To account for uncontrolled differences in how fitness was measured between studies, we allowed mean fitness effects to randomly vary between studies by including random intercepts for each study in the mixed model. A similar mixed linear model was used to model the relationship between explanatory features and the magnitude of pleiotropy. We used logistic regression to test for significant associations between explanatory features and the sign of pleiotropy (positive or antagonistic). For the latter, we used standard as opposed to mixed logistic models as the mixed models did not converge. Finally, we tested whether the observed distribution of pleiotropic effects grouped by their sign (+/+, +/-, -/-) was different from what would be expected given the observed distribution of fitness effects in single hosts using a Pearson’s chi-squared test. The chi-square test was performed in two ways to ensure the random ordering of hosts as Host_1_ or Host_2_ did not change the significance of the results. First, all mutations with antagonistic fitness effects across hosts were summed together and then evenly divided between the +/- and -/+ entries in the contingency table constructed for the chi-square test (as in Table 3). Second, to ensure that a single random ordering did not bias the test results, we randomly permuted the ordering of hosts and reperformed the chi-square test 1,000 times.

### Sensitivity analysis

Many studies only report mutational fitness effects as point estimates or mean effects, making it difficult to disentangle true variability in fitness effects from measurement error. Because a large fraction of mutations also have small fitness effects close to zero (neutral), even low levels of error could result in the sign of the fitness effect being reversed, causing us to misinfer the sign of pleiotropy as a result. We therefore sought to understand how different levels of error in measured fitness effects would impact the distribution of pleiotropic fitness effects between hosts. To this end, we simulated higher levels of measurement error by introducing additional noise or error in the reported fitness effects. Specifically, for each mutation in each host, we included an additional error term randomly drawn from a Normal distribution centered at the reported fitness value with a standard deviation σ that we varied to emulate increasing levels of error. This error term was drawn independently from the same distribution in each host, such that the errors were uncorrelated between pairs of hosts. We note that assuming independence between hosts is conservative with respect to any hypothesis about the sign of pleiotropy, whereas assuming fitness effects are negatively or positively correlated between hosts would skew the distribution towards antagonistic or positive pleiotropic effects. After each simulation, summary statistics describing the distribution of pleiotropic effects were recomputed and we report the mean statistic computed from 1,000 simulations run at each level of error (σ).

## Results

We identified 26 studies in which the fitness effects of individual viral mutations were experimentally determined in at least two host environments (Supplemental Table S1). In all, our data set includes fitness effects for 153 mutations in 20 unique viruses and 34 unique hosts. The resulting data set allows us to make 681 pairwise comparisons of fitness effects between hosts in order to quantify the overall distribution of pleiotropic effects. In order to standardize fitness effects across studies, we quantify the relative fitness effect of a mutation in each host as ω = *log_2_(w_mut_ / w_ref_)*, where *w_mut_*and *w_ref_* are the fitness of the mutant and wildtype virus, respectively. A neutral mutation therefore has a relative fitness effect ω = 0, whereas beneficial mutations have fitness effects ω > 0 and deleterious mutations have ω < 0. On this log scale, a mutation that increases fitness two-fold will have ω = 1.

### Distribution of mutational fitness effects in single hosts

As expected from previous work characterizing the distribution of fitness effects (DFE) of viral mutations in single hosts (Domingo-Calap et al., 2009; Eyre-Walker & Keightley, 2007; Sanjuán et al., 2004), most non-lethal mutations are nearly neutral with a prominent peak at ω = 0, but the distribution is skewed towards deleterious effects (Figure 1; Fisher-Pearson skewness = -0.25). Out of 457 total fitness effects, 66.8% are deleterious in individual hosts. Averaged over all hosts, the mean fitness effect is -0.205 (95% Confidence Interval (CI): -0.306:-0.102), which translates to a 13.6% decrease in fitness on a linear scale.

**Figure 1:**
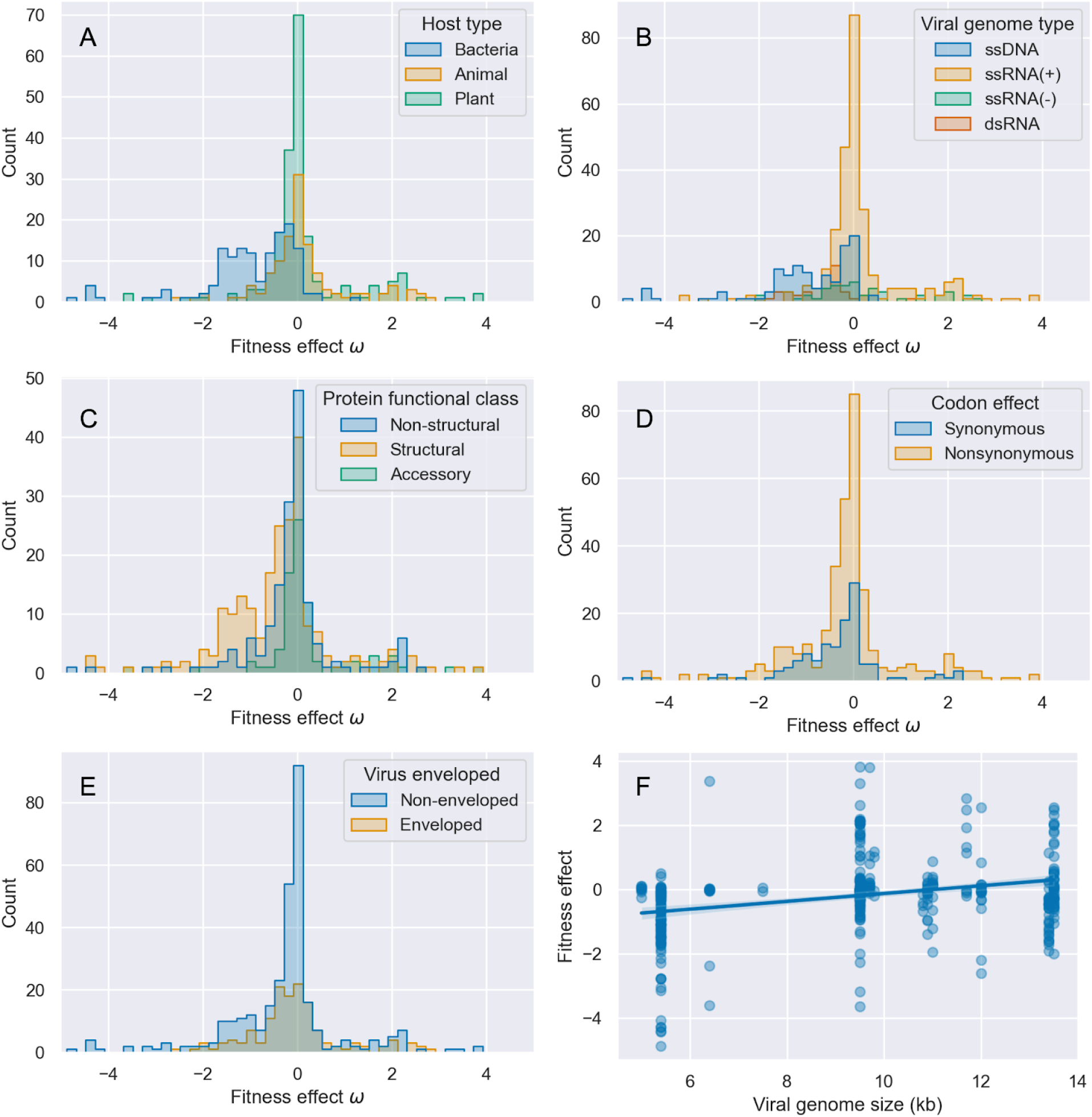
Distribution of mutational fitness effects in single hosts. Fitness effects ω are plotted on a log_2_ scale. The fitness effects are grouped by: (A) host type, (B) viral genome type, (C) viral protein functional class, (D) codon effect, and (E) viral envelope status. (F) The relationship between viral genome size and mutational fitness effects.

We further explored whether any features explain the observed variability in fitness effects in single hosts (Figure 1; Table 1). To account for potential confounding effects among features as well as uncontrolled differences between studies, we estimated the impact of each feature on mutational fitness effects using a mixed effect linear model. Explanatory features were treated as fixed effects while allowing mean fitness effects (intercepts) to randomly vary by study. Only two features were estimated to significantly impact mutational fitness effects. First, mutations in bacteriophages are on average more deleterious than mutations in plant and animal viruses (Figure 1A). As a result, mutations in single-stranded DNA viruses, which are largely bacteriophages, tend to be more deleterious than mutations in positive (+) or negative (-) single-stranded RNA viruses, but not significantly so (Figure 1B).Mutations tend to be slightly less deleterious in viruses with larger genomes (Figure 1F). On the other hand, fitness effects did not differ significantly between mutations in structural proteins versus mutations in non-structural and accessory proteins (Figure 1C), nor did they significantly differ between nonsynonymous and synonymous mutations (Figure 1D), nor between enveloped and non-enveloped viruses (Figure 1E).The intraclass correlation coefficient (ICC) among studies is 0.341, indicating that a substantial amount of variance in fitness effects can be explained by between study variation in reported fitness effects.

**Table 1:** Mutational fitness effects in single hosts grouped by viral features. Regression coefficients, P-values and 95% CIs are estimated under a mixed effect linear model.

| Variable | N | Group mean | Estimated coefficient | P-value | 95% CI |
| --- | --- | --- | --- | --- | --- |
| Intercept | 457 | - | -3.639 | 0.65 | -8.043 : 0.766 |
| Synonymous | 121 | -0.37 | - | - | - |
| Nonsynonymous | 336 | -0.14 | -0.063 | 0.644 | -0.329 : 0.204 |
| Accessory | 83 | 0.25 | - | - | - |
| Non-structural | 156 | -0.13 | -0.19 | 0.225 | -0.497 : 0.117 |
| Structural | 218 | -0.42 | 0.368 | 0.108 | -0.081 : 0.816 |
| dsRNA | 36 | -0.69 | - | - | - |
| ssDNA | 112 | -1.01 | 0.958 | 0.462 | -1.593 : 3.509 |
| (+)ssRNA | 243 | 0.12 | 0.14 | 0.889 | -1.834 : 2.115 |
| (-)ssRNA | 46 | 0.26 | -1.299 | 0.211 | -3.336 : 0.738 |
| Animal host | 119 | 0.13 | - | - | - |
| **Bacteria host** | 139 | -0.98 | -1.761 | **0.024** | -3.288 : -0.235 |
| Plant host | 199 | 0.14 | -0.659 | 0.478 | -2.480 : 1.162 |
| Enveloped | 144 | -0.06 | - | - | - |
| Non-enveloped | 313 | -0.27 | 1.289 | 0.236 | -0.841 : 3.419 |
| **Viral genome size** | 457 | - | 0.329 | **0.032** | 0.028 : 0.630 |

We also fit a second regression model that included viral family as the only fixed effect and study as a random effect. This allowed us to explore whether fitness effects vary between viral taxonomic groups independently of their host or viral genome type while still accounting for differences between studies. While mean fitness effects varied considerably between viral families, no family had a significant impact on mutational fitness effects (Supplemental Table S3). Together, these results suggest that aside from mutations being slightly more deleterious on average in bacteriophages, the DFE is roughly similar across different viruses and hosts.

### Magnitude of pleiotropy

We compared the fitness effects of each mutation between every unique pair of hosts for which fitness data was available, allowing us to characterize the overall distribution of pleiotropic fitness effects. We first consider the magnitude of pleiotropy, where the magnitude is defined as the absolute difference in the fitness effect of a mutation between two hosts, such that the magnitude is larger the more different the fitness effects are between hosts, regardless of the sign of the fitness effects.

Because we expect fitness effects to differ more between more phylogenetically divergent hosts (Longdon et al., 2014, 2018), we first considered the relationship between host divergence times and the magnitude of pleiotropy (Figure 2). When considered among all pairs of hosts, divergence times do not have a significant effect on the magnitude of pleiotropy (β*_div_* = -0.005; 95% CI = -0.052 : 0.042; p=0.832). However, when we consider interactions between host type and divergence times, the magnitude of pleiotropy increases strongly and significantly with divergence times for plants (β*_div X plant_* = 0.985: 95% CI = 0.786 : 1.184; p<0.001), weakly but not significantly for bacteria (β*_div X bacteria_* = 0.109; 95% CI = -0.013 : 0.23; p=0.079), but not for animals (β*_div X animal_*= 0.016, 95% CI = -0.029 : 0.06; p=0.49).

**Figure 2:**
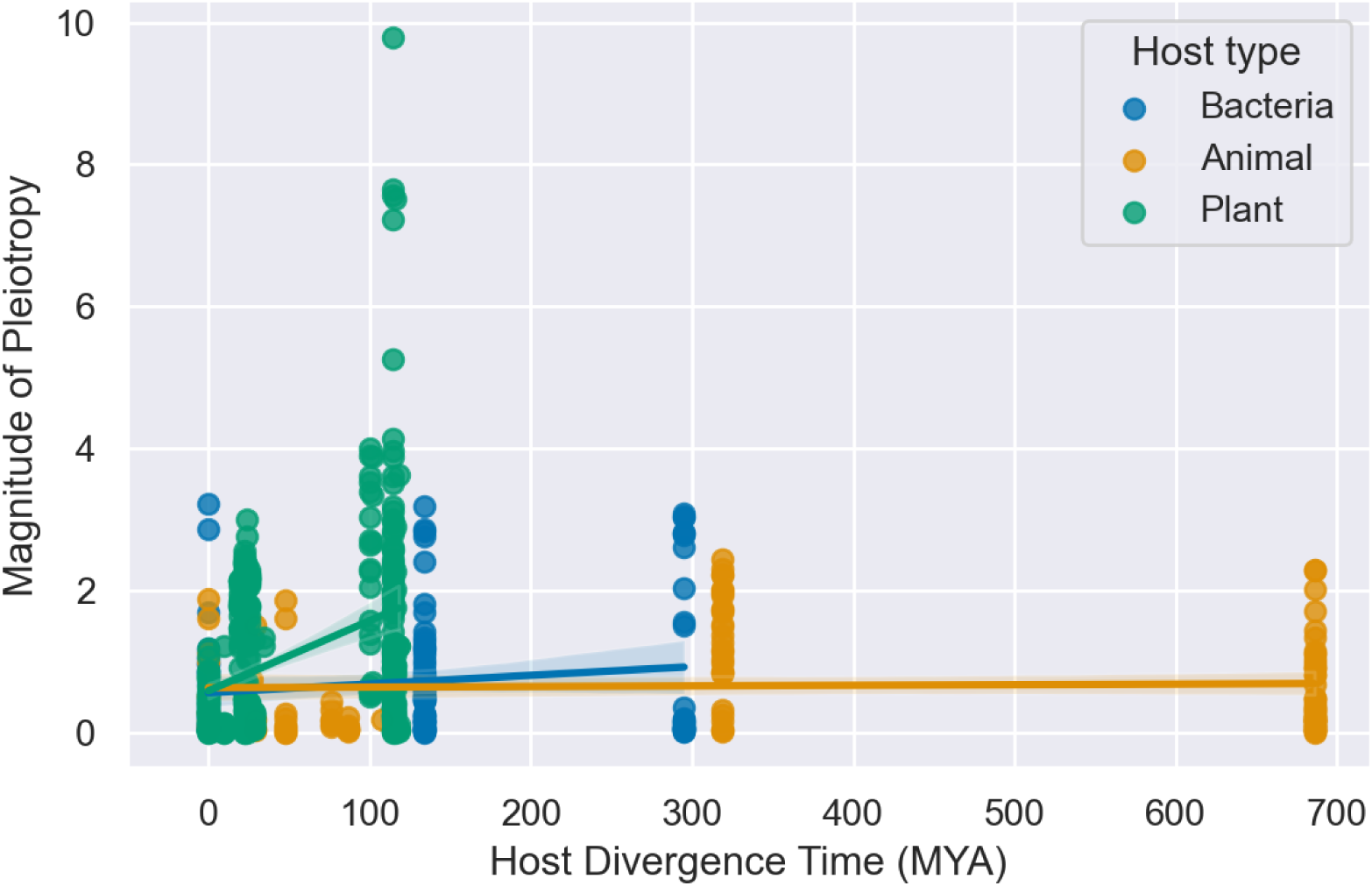
Magnitude of pleiotropic fitness effects between pairs of hosts regressed against host divergence times in millions of years. Colored lines show the estimated regression coefficients (slope) for each host type.

We also considered how the magnitude of pleiotropy depends on several other explanatory features (Figure 3, Table 2). After controlling for confounding effects and uncontrolled differences between studies using a mixed effect model, no feature was estimated to have a significant effect on the magnitude of pleiotropy. However, when we included interactions between host type and host divergence times in the model, divergence times between plant hosts have a strong and significant effect on the magnitude of pleiotropy (Table 2). Thus of all the features we considered only divergence times between plant hosts strongly predict the magnitude of pleiotropy. Under the mixed effect model, the intraclass correlation coefficient among studies is low (ICC=0.055), indicating the magnitude of pleiotropic effects does not significantly vary between studies.

**Figure 3:**
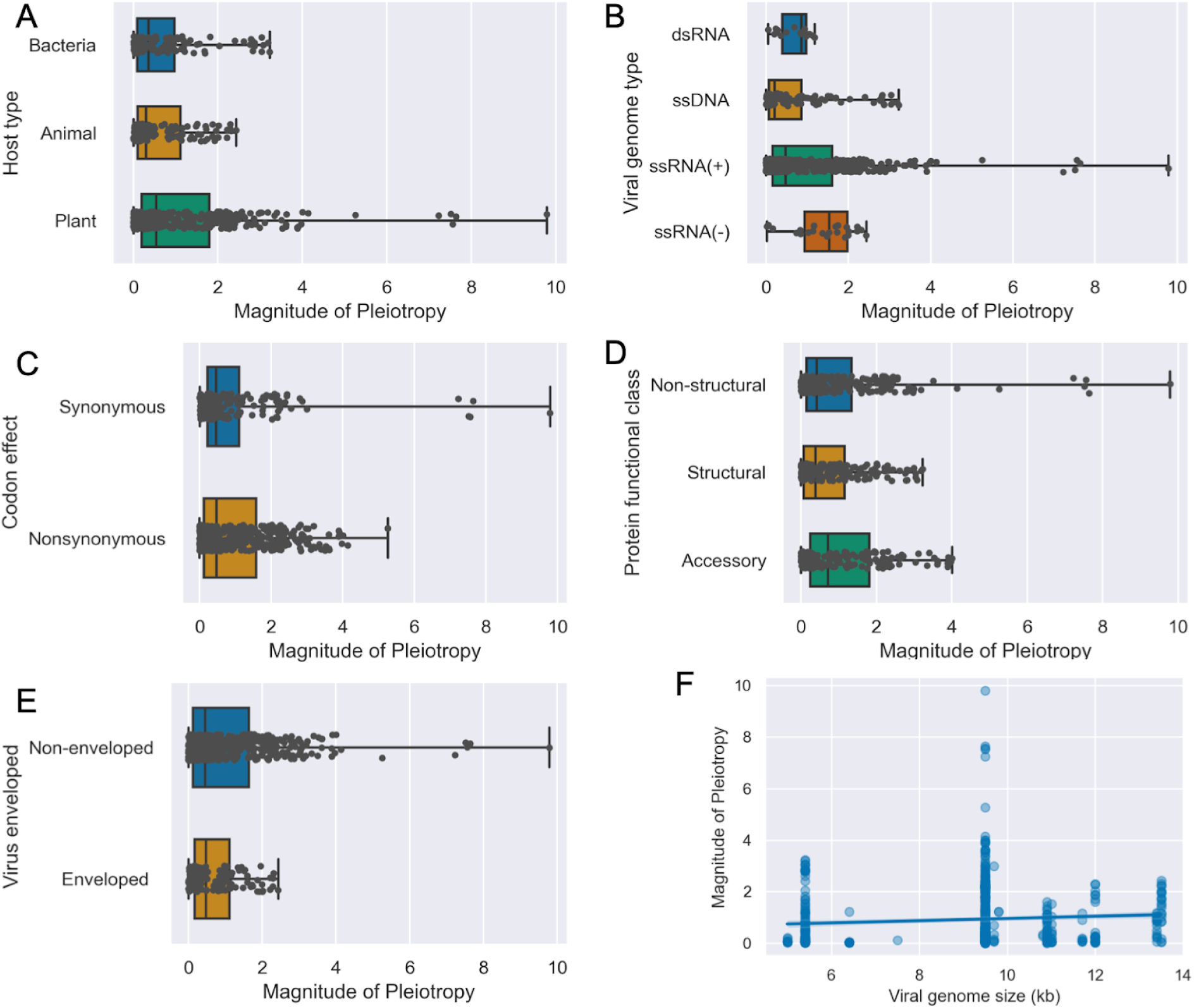
Magnitude of pleiotropic fitness effects computed as the absolute difference in fitness effects between hosts. Fitness effects in each host were first log_2_ transformed, such that a magnitude of one reflects a 2X difference in fitness effects between hosts. Box plots show the median and quartiles of each categorical group while whiskers show the full range of observed values.

**Table 2:** Magnitude of pleiotropic fitness effects grouped by viral features. Regression coefficients, P-values and 95% CIs are estimated under a mixed effect linear model.

| Variable | N | Group mean | Estimated coefficient | P-value | 95% CI |
| --- | --- | --- | --- | --- | --- |
| Intercept | 681 | - | 0.392 | 0.535 | -0.845 : 1.628 |
| Synonymous | 183 | 0.92 | - | - | - |
| Nonsynonymous | 498 | 0.93 | -0.148 | 0.193 | -0.370 : 0.075 |
| Accessory | 180 | 1.1 | - | - | - |
| Non-structural | 299 | 0.92 | 0.101 | 0.364 | -0.117 : 0.319 |
| Structural | 202 | 0.76 | 0.254 | 0.191 | -0.127 : 0.635 |
| dsRNA | 18 | 0.73 | - | - | - |
| ssDNA | 124 | 0.67 | 0.012 | 0.992 | -2.257 : 2.280 |
| (+)ssRNA | 516 | 0.96 | 0.021 | 0.974 | -1.226 : 1.268 |
| (-)ssRNA | 23 | 1.42 | 0.804 | 0.248 | -0.560 : 2.168 |
| Enveloped | 129 | 0.72 | - | - | - |
| Non-enveloped | 552 | 0.97 | -0.28 | 0.795 | -2.392 : 1.832 |
| Animal host | 121 | 0.66 | - | - | - |
| Bacteria host | 133 | 0.87 | 0.228 | 0.658 | -0.783 : 1.240 |
| Plant host | 427 | 1.17 | 0.274 | 0.798 | -1.827 : 2.374 |
| Divergence time | 681 | - | 0.02 | 0.58 | -0.052 : 0.093 |
| Divergence x<br>Animal host | 121 | - | - | - | - |
| Divergence x<br>Bacteria host | 133 | - | 0.151 | 0.158 | -0.059 : 0.361 |
| **Divergence x**<br>**Plant host** | 427 | - | 0.87 | **<0.001** | 0.616 : 1.123 |

### Sign of pleiotropy

Across all pairwise comparisons of fitness effects, positive pleiotropy is slightly more frequent (391/681 = 57.4%) than antagonistic pleiotropy. Likewise, mutational fitness effects are overall positively correlated between hosts (Pearson correlation coefficient ρ = 0.32). However, our definition of positive pleiotropy includes mutations that are both unconditionally beneficial and unconditionally deleterious. If we only consider pairwise comparisons with positive pleiotropic effects, the majority are unconditionally deleterious (297/391 = 75.9%). Nevertheless, 13.8% of all pairwise comparisons are unconditionally beneficial, suggesting that mutations that increase fitness in more than one host are not uncommon.

We next tested whether the observed frequency of each type of sign pleiotropy (+/+, +/-, -/-) differed from what would be expected based on the number of beneficial and deleterious mutations observed in single hosts (Table 3). A Pearson chi-square test indicates the observed frequencies are consistent with what would be expected by chance if the pairwise fitness effects were randomly drawn from the distribution of fitness effects in single hosts (χ^2^ = 2.62, p-value = 0.106). In other words, antagonistic and positive pleiotropy arise at roughly the frequency we would expect given the relative number of beneficial versus deleterious mutations observed in individual hosts. To further ensure these results were not driven by the random ordering of hosts, we randomly permuted the ordering of hosts and repeated the chi-square test 1,000 times. The chi-square test was always nonsignificant, with p-values ranging from 0.104 to 0.132. Given a mutation is beneficial in one host, it might be expected that the likelihood of the mutation being deleterious in another host would increase with the size of the beneficial fitness effect in the first host. However, the odds of a mutation being beneficial in both hosts actually increases with the size of the beneficial fitness effect in the first host (β = 1.17; 95% CI = 1.07 : 1.29; p=0.001).

**Table 3:** Contingency table with the observed versus expected number of mutations in each pleiotropic class. Mutations with antagonistic pleiotropic effects were evenly divided between the +/- and -/+ cells as the ordering of hosts is arbitrary.

|  |  | Host 2 Fitness Effect |  |
| --- | --- | --- | --- |
|  |  | Negative (-) | Positive (+) |
| Host 1 Fitness Effect | Positive (+) | Obsv = 145 (21.3%)<br>Exp = 159.5 (23.3%) | Obsv = 94 (13.8%)<br>Exp = 78 (11.5%) |
|  | Negative (-) | Obsv = 297 (43.6%)<br>Exp = 282 (41.4%) | Obsv = 145 (21.3%)<br>Exp = 159.5 (23.3%) |

We further explored whether certain types of mutations were more likely to have positive rather than antagonistic pleiotropic effects using logistic regression (Table 4). The odds of a mutation having positive pleiotropic effects are higher in ssDNA viruses, and by extension bacteriophages, but these mutations were nearly always unconditionally deleterious (Figure 4A-B). Synonymous mutations are no more likely to have positive pleiotropic effects than nonsynonymous mutations (Figure 4C). Although not statistically significant, mutations in structural proteins are more likely to have positive pleiotropic effects, but again most mutations in structural proteins are unconditionally deleterious (Figure 2D). Finally, mutations in non-enveloped viruses are significantly less likely to exhibit positive pleiotropy than enveloped viruses (Figure 4E).

**Figure 4:**
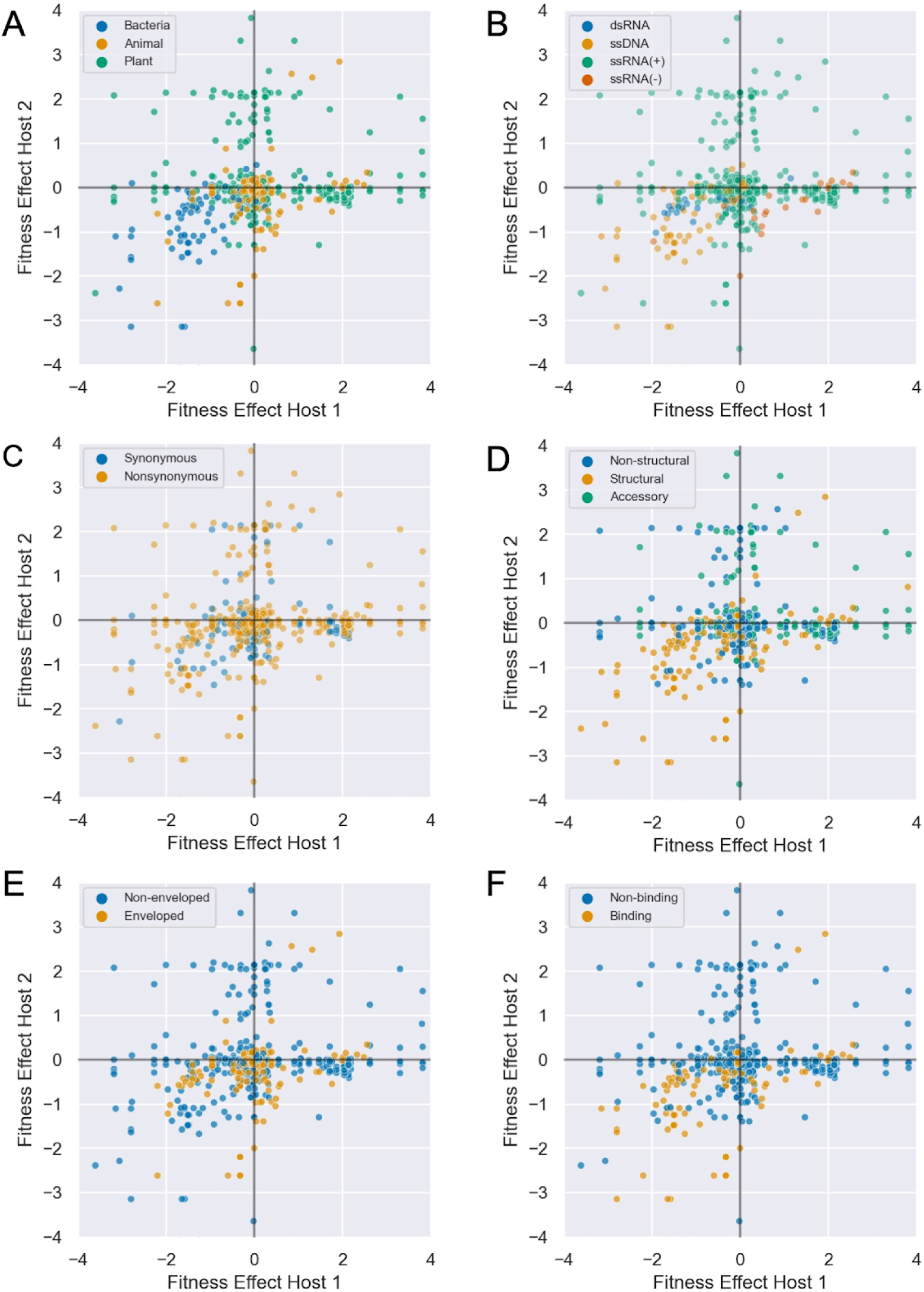
Joint distribution of mutational fitness effects between pairs of hosts. The underlying distribution is identical in all plots but mutations are colored by: (A) host type, (B) viral genome type, (C) codon effect, (D) protein functional class, (E) viral envelope status and (F) whether the mutation occurs in a protein involved in host cell binding. Fitness effects ω in each host are log_2_ transformed.

**Table 4:** Odds of a mutation having a positive pleiotropic effect. Odds ratios, p-values and 95% CI are estimated under a multivariate logistic regression model.

| Variable | Frequency | Odds ratio | P-value | 95% CI |
| --- | --- | --- | --- | --- |
| Intercept | - | 10.83 | 0.38 | - |
| Synonymous | 92/183 | - | - | - |
| Nonsynonymous | 284/498 | 0.82 | 0.36 | 0.54 : 1.25 |
| Accessory | 81/180 | - | - | - |
| Non-structural | 138/299 | 0.71 | 0.13 | 0.46 : 1.1 |
| Structural | 157/202 | 1.22 | 0.55 | 0.64 : 2.33 |
| dsRNA | 16/18 | - | - | - |
| ssDNA | 114/124 | 3.32 | 0.4 | 0.20 : 55.4 |
| (+)ssRNA | 236/516 | 0.29 | 0.19 | 0.04 : 1.80 |
| **(-)ssRNA** | 10/23 | 0.1 | **0.006** | 0.02 : 0.52 |
| Enveloped | 81/129 | - | - | - |
| **Non-enveloped** | 295/552 | 0.38 | **0.03** | 0.15 : 0.92 |
| Viral genome size (kb) | - | 0.98 | 0.9 | 0.67 : 1.41 |

Among mutations exhibiting positive pleiotropy, we further examined whether any features increase the odds of mutations being unconditionally beneficial rather than deleterious (Table 5). The odds of mutation being unconditionally beneficial are slightly higher for nonsynonymous than synonymous mutations. Mutations in (-)ssRNA viruses also have significantly higher odds of being unconditionally beneficial than in viruses with other genome types, but this was based on a small number of observations. On the other hand, mutations in both structural and non-structural proteins are more likely to be unconditionally deleterious than mutations in accessory proteins, although only significantly for non-structural proteins. To better understand what types of proteins tend to increase the odds of unconditional deleterious effects, we further divided proteins by their more specific molecular function (Supp. Figure S1) and whether or not the protein is involved in host cell-binding (Figure 2F). While no clear pattern emerged in terms of protein molecular function, mutations in structural proteins like glycoproteins that bind to host cells are significantly more likely to be unconditionally deleterious than mutations in proteins that do not bind to host cells.

**Table 5:** Odds of a mutation being unconditionally beneficial versus deleterious. Odds ratios, p-values and 95% CI are estimated under a multivariate logistic regression model.

| Variable | Frequency | Odds ratio | P-value | 95% CI |
| --- | --- | --- | --- | --- |
| Intercept | - | 0.05 | 0.56 | - |
| Synonymous | 11/92 | - | - | - |
| **Nonsynonymous** | 68/284 | 2.21 | **0.048** | 1.01 : 4.84 |
| Accessory | 31/81 | - | - | - |
| **Non-structural** | 32/138 | 0.34 | **0.009** | 0.15 : 0.77 |
| Structural | 16/157 | 0.44 | 0.203 | 0.12 : 1.56 |
| dsRNA | 1/16 | - | - | - |
| ssDNA | 7/114 | 5.81 | 0.47 | 0.05 : 710.5 |
| (+)ssRNA | 65/236 | 2.43 | 0.51 | 0.16 : 34.85 |
| **(-)ssRNA** | 6/10 | 19.18 | **0.016** | 1.73 : 212.1 |
| Enveloped | 27/81 | - | - | - |
| Non-enveloped | 52/295 | 0.34 | 0.18 | 0.07 : 1.65 |
| Non-host cell binding | 69/250 | - | - | - |
| **Host cell binding** | 10/126 | 0.08 | **0.002** | 0.02 : 0.41 |
| Viral genome size (kb) | - | 1.22 | 0.58 | 0.60 : 2.49 |

### Sensitivity analysis

Because mutational fitness effects are often reported without any measure of error or uncertainty, we performed a sensitivity analysis to see how robust our main results are to different levels of simulated measurement error in the reported fitness effects (Supp. Table S4). With increasing levels of simulated error, the positive correlation in fitness effects between hosts weakens, suggesting that in the absence of error, the true fitness effects would have been more positively correlated between hosts. Likewise, increasing levels of error causes the fraction of mutations with antagonistic effects between hosts to increase relative to the fraction with concordant effects, suggesting that positive pleiotropy may be even more common in the absence of error. At the same time, increasing levels of error causes the number of mutations with unconditionally beneficial effects to increase relative to the number with unconditionally deleterious effects. This suggests that the true fraction of unconditionally beneficial mutations may be smaller than the 13.8% reported above. Finally, we reperformed the chi-square test for the independence of the sign of fitness effects between hosts at different levels of error. The χ^2^ statistic shows a non-monotonic relationship with the level of error, first increasing at low levels and then decreasing at high levels of error. This suggests that fitness effects would become increasingly independent between hosts at higher levels of error. While this also suggests that fitness effects would become increasingly independent in the absence of error, we cautiously interpret these results to suggest that our finding about the independence of fitness effects between hosts is at least robust to measurement error. Taken together, the sensitivity analysis indicates that in the absence of error the relative frequency of positive pleiotropy relative to antagonistic pleiotropy would have increased.

## Discussion

Viral host ranges are widely believed to be constrained by fitness tradeoffs arising from antagonistic pleiotropy (Bedhomme et al., 2015; Duffy et al., 2000; Elena et al., 2009). But to our knowledge, no previous work has systematically surveyed the fitness effects of mutations in different hosts to quantify the relative frequency of antagonistic versus positive pleiotropy. Within individual hosts, our meta-analysis confirmed that the distribution of fitness effects (DFE) is skewed towards deleterious mutations with individual mutations reducing fitness by 13.6% on average. This is remarkably similar to previous estimates which estimated mutations reduce fitness 10.3 - 13.2% on average (Sanjuán, 2010). Considering the pairwise or joint DFE between hosts, fitness effects are fairly strongly positively correlated between hosts and positive pleiotropy is actually slightly more common than antagonistic pleiotropy. While antagonistic pleiotropy may nevertheless constrain the ability of viruses to adapt to certain hosts, our study calls into question whether antagonistic pleiotropy by itself is frequent and strong enough to constrain viral host ranges more generally.

Perhaps our most surprising result is that antagonistic pleiotropy is no more frequent than would be expected by chance given the frequency of beneficial and deleterious mutations in individual hosts. Because the DFE is skewed towards deleterious mutations in individual hosts, we expect a majority of mutations to be unconditionally deleterious between hosts, a moderate fraction to have antagonistic effects and very few to be unconditionally beneficial in both hosts. The observed distribution of pleiotropic fitness effects is remarkably similar to this expectation, suggesting that the sign of a mutation’s fitness effect in one host is essentially independent of its sign in another. This is consistent with earlier work on individual viruses showing the fitness effects of new mutations are positively correlated between hosts overall but the sign of fitness effects are highly unpredictable between hosts (Lalić et al., 2011; Vale et al., 2012). This may also explain why we found few features of mutations, viruses or hosts that could reliably predict whether or not mutations will have antagonistic effects.

Given that there appears to be no natural tendency towards mutations having antagonistic effects between hosts, antagonistic pleiotropy may be less ubiquitous than previously thought. This may at first seem surprising; afterall viruses are obligate cellular parasites requiring many finely tuned molecular interactions between viral and host factors. However, even if fitness effects are host-specific, these effects may only vary by degree between hosts, impacting the magnitude of pleiotropy but not the sign. Many viral proteins may also interact with similar host factors in different hosts, especially if the hosts are closely related (Damas et al., 2020; Long et al., 2018). As might be expected based on these considerations, we found that fitness effects are more similar between pairs of more phylogenetically related hosts, at least in bacteria and plant hosts. Furthermore, mutations in certain viral proteins may even be expected to have concordant fitness effects between hosts. For example, improving the processivity of RNA polymerases or the stability of structural proteins may increase fitness across a wide range of hosts (Duffy et al., 2000; Manz et al., 2013). In other microbial systems, positive pleiotropy likewise appears more common than previously thought, including in bacteria growing on different carbon sources (Leiby & Marx, 2014; Ostrowski et al., 2005; Sane et al., 2018) and in different antibiotic environments (Ardell & Kryazhimskiy, 2021; Chevereau et al., 2015; Reding-Roman et al., 2017). Thus, rather than being viewed as surprising, reports of positive pleiotropy should often be expected.

The more pertinent question for long-term viral evolution may be whether the observed distribution of pleiotropic effects can prevent the evolution of generalist viruses adapted to multiple hosts. On longer timescales, purifying selection will remove most unconditionally deleterious mutations, such that it is ultimately the availability of beneficial mutations on the timescales relevant to adaptation in each host that determines whether generalists evolve. Since it is commonly observed that viruses do adapt to multiple hosts when rapidly passaged between alternate hosts (Bedhomme et al., 2012; Coffey & Vignuzzi, 2010; Kassen, 2002; Smith-Tsurkan et al., 2010; Weaver et al., 1999), it appears that the supply of unconditional beneficial mutations is often sufficient for generalists to evolve. Indeed, our meta-analysis revealed that unconditionally beneficial mutations are not especially rare (13.8% of all pairwise comparisons). However when viruses infrequently encounter an alternate host, antagonistic pleiotropy may be sufficiently common to constrain the evolution of generalists (Bono et al., 2017; Visher & Boots, 2020). In this case, beneficial mutations will be selected in the more frequently encountered host irrespective of their fitness effects in the alternate host. Thus, even if antagonistic pleiotropy is no more common than expected by chance, mutations fixed in the more frequent host will on average be deleterious in the rare host. The resulting accumulation of deleterious mutations can lead to the evolution of specialists adapted only to the more common host, as proposed under the mutation accumulation hypothesis (Kawecki, 1994; Whitlock, 1996). Thus, whether or not antagonistic pleiotropy is sufficient to constrain the evolution of generalist viruses may strongly depend on the frequency at which different hosts are encountered.

Interestingly, many studies report both cases where antagonistic pleiotropy constrains and positive pleiotropy promotes adaptation to different hosts, often simultaneously in the same host-virus pair (Duffy et al., 2000; Moreno-Pérez et al., 2016; Novella et al., 2011; Remold et al., 2008; Visher et al., 2022). Theory should therefore be able to explain why pleiotropy sometimes constrains adaptation and yet other times promotes the evolution of multi-host generalists. We propose a simple conceptual model for how pleiotropy may either constrain or promote adaptation based on general observations of the DFE that should apply across many host-pathosystems (Figure 5). Similar to other recent work, we view fitness tradeoffs along a continuum from hard to soft based on how easy it would be for evolution to resolve or overcome a tradeoff (Ardell & Kryazhimskiy, 2021; Tikhonov et al., 2020). Hard or “rigid” tradeoffs reflect absolute constraints arising from conflicting structural or biophysical requirements in different hosts. In this case, viral populations may be evolving along a Pareto optimality front where fitness cannot be increased in one host without compromising fitness in an alternate host (Li et al., 2019; Shoval et al., 2012). At such a Pareto front, the biophysical constraints underlying the tradeoff may skew the distribution of fitness effects towards mutations with antagonistic and unconditionally deleterious effects (the orange dots in Figure 5A). Selection is therefore unable to resolve such tradeoffs because mutations that simultaneously increase fitness in both hosts are not accessible.

**Figure 5:**
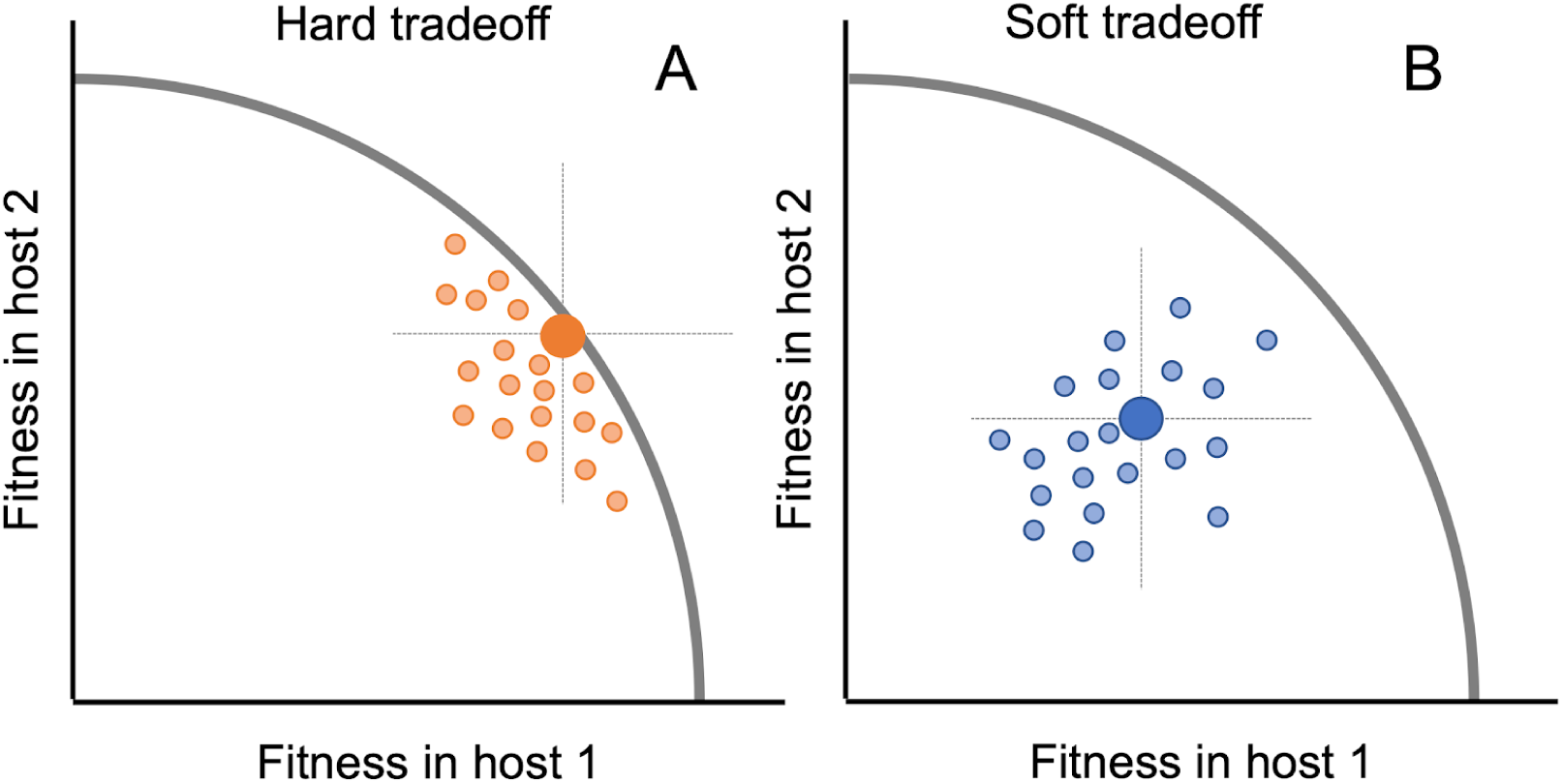
Conceptual model illustrating how the distribution of pleiotropic effects can lead to both hard and soft fitness tradeoffs between hosts. Small dots reflect the joint distribution of fitness effects (DFE) of mutations in each host. Crosshairs divide the joint DFE into four quadrants based on whether mutations have beneficial or deleterious effects in each host relative to a reference genotype (large dots). (A) Hard tradeoffs arise when absolute biophysical constraints, represented as a hypothetical Pareto front (curved line), prevent fitness from improving in one host without declining in the other host. In this case, the joint DFE (orange dots) is shifted towards mutations with antagonistic or unconditionally deleterious effects. (B) Soft tradeoffs arise when a population lies behind a Pareto front, shifting the joint DFE (blue dots) towards unconditionally beneficial mutations.

In contrast to hard tradeoffs, soft or “plastic” tradeoffs can easily be overcome by evolution. Soft tradeoffs can arise when a population lies behind a hypothetical Pareto front, possibly due to a lack of prior selection in one or both environments. In this case, the distribution of fitness effects may be shifted towards positive pleiotropy such that unconditionally beneficial mutations are accessible, allowing a population to rapidly adapt to both hosts without paying any apparent cost (Figure 5B). This provides a general mechanism whereby viruses could overcome fitness tradeoffs that is strongly supported by our knowledge of the DFE. In populations that are poorly adapted to their environment, the entire DFE shifts towards beneficial mutations which increase in both frequency and effect sizes (Chou et al., 2011; Eyre-Walker & Keightley, 2007; Kryazhimskiy et al., 2014; MacLean et al., 2010). Likewise, if we consider two environments, the joint DFE may shift towards unconditionally beneficial mutations as a result of the marginal DFE in each host shifting towards beneficial effects (Wei & Zhang, 2019).

The overall distribution of pleiotropic fitness effect we observe here, where fitness effects are overall positively correlated between hosts with many unconditionally beneficial mutations, implies that most fitness tradeoffs between hosts are soft and could thus be easily overcome. However, one caveat to this conclusion is that, because our meta-analysis combines fitness effects from many different viruses and hosts, it may not be representative of the DFE in any virus-host pair *per se*, but rather reflect a mixture of the different distributions shown in Figure 5. Thus, the general occurrence of unconditionally beneficial mutations does not necessarily imply that beneficial mutations will be accessible in any specific host-virus system. Additionally, individual studies often measure different components of viral fitness and the measured component may differ from those components underlying fitness tradeoffs in nature. For example, studies of animal viruses typically measure fitness in cell culture such that fitness reflects cellular infection rates, whereas studies of plant viruses tend to measure fitness in whole plants, such that fitness reflects both replication and movement within a systemically infected host. While we tried to mitigate these differences by only comparing mutational fitness effects pairwise between hosts measured in the same study, the relative frequency of antagonistic versus positive pleiotropy may vary based on what component of fitness is being considered..

Nevertheless, we conjecture that the overall distribution of pleiotropic fitness effects observed here likely is representative of the situation many viruses experience as they adapt to novel hosts. To viruses, novel hosts represent low-quality, stressful environments which tend to skew the DFE towards beneficial mutations (Martin & Lenormand, 2006; Vale et al., 2012). Many of these mutations may be beneficial across multiple hosts, allowing for fitness trade ups rather than tradeoffs. On the other hand, hard tradeoffs may only arise in situations where historical selection pressures have nearly optimized fitness in both hosts, as seen in some vector-borne viruses (Coffey et al., 2008; Cooper & Scott, 2001; Greene et al., 2005). Unfortunately, existing data make it difficult to explicitly test these theories because mutational fitness effects are generally only considered in a single genetic background and it is often unclear to what extent these reference genotypes were previously adapted to the experimental environment (Ferris et al., 2007; Vale et al., 2012). With additional information about where a viral population lies in its overall fitness landscape, it may be possible to predict the distribution of pleiotropic effects, and thus whether tradeoffs are likely to be hard or soft.

Finally, while theoretical models of fitness tradeoffs generally try to explain the evolutionary success of either specialists or generalists, in nature both specialist and generalist viruses co-exist and host ranges vary in breadth from single hosts to thousands of hosts spanning multiple kingdoms (Andika et al., 2017; Parrella et al., 2003; Wilson & Yoshimura, 1994). This variability in host range breadth is only explicable if we consider the different selection pressures viruses experience alongside genetic factors like the DFE. For example, a plant virus vectored by a herbivorous insect may repeatedly experience selection to adapt to any plant host on which its vector next alights (Lefeuvre et al., 2019). Meanwhile, a phage specialized on an abundant bacterial host may experience effectively no selection to adapt to a less common host. Thus, while the widespread prevalence of positive pleiotropy suggests many tradeoffs may be soft, the right historical selection pressures still need to be in place for viruses to expand their host range. A more fully developed theory of viral host range evolution would therefore consider how both biophysical constraints as well as past and current selection pressures shape the distribution of pleiotropic fitness effects and thereby the strength of fitness tradeoffs between hosts.

## Data and code availability

Our full dataset containing all extracted fitness effects and metadata is available at https://github.com/davidrasm/PleiotropyMetaAnalysis.

