## Supplemental Material for "Quantifying the strength of viral fitness tradeoffs between hosts: A meta-analysis of pleiotropic fitness effects"

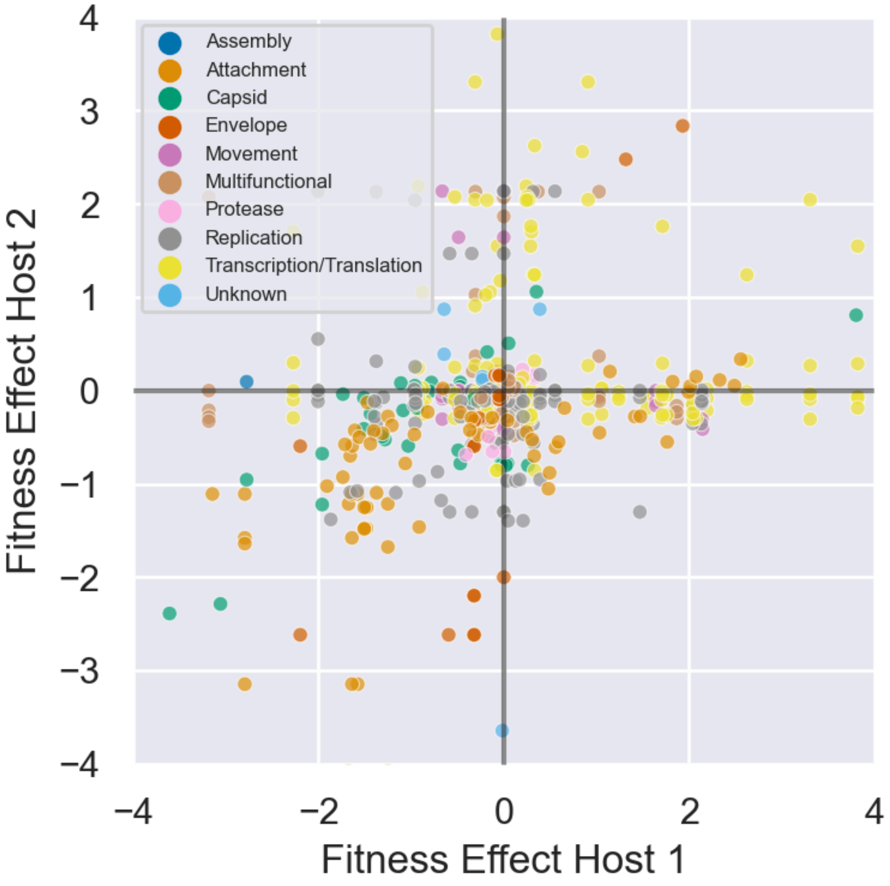


**Figure S1:** Joint distribution of mutational fitness effects between pairs of hosts colored by the molecular function of the protein in which each mutation occurs.

**Table S1**. List of the 26 published journal articles used in our meta-analysis.

| Authors | Year | Journal | Title | Virus | Hosts | Data source | Quantification method |
|---|---|---|---|---|---|---|---|
| Pepin et al. | 2006 | Genetics | Variable Pleiotropic Effects From Mutations at the Same Locus Hamper Prediction of Fitness From a Fitness Component | PhiX174 | *Escherichia coli* C (Ec C); *Escherichia coli* (Ec waaL); *Shigella sonnei* (Ss); *Salmonella enterica serovar typhimurium* (St rfb); *Salmonella enterica* (St GalE); *Escherichia coli* (Ec waaV) | Fig.2 and Fig.3 | Growth rate (virus titer); Webplot digitizer |
| Vale et al. | 2012 | Evolution | The Distribution of Mutational Fitness Effects of Phage φX174 on Different Hosts | PhiX174 | *Escherichia coli; Salmonella typhimurium* | Fig. 1 | Fitness difference conversion to ratios; Webplot digitizer |
| Ferris et al. | 2007 | Genetics | High Frequency of Mutations That Expand the Host Range of an RNA Virus | Phi6 | *Pseudomonas syringae* (pathovar phaseolicola); *Pseudomonas syringae* (pathovar japonica) | Contacted author and requested data | Plaque size fitness assay |
| Hanley et al. | 2003 | Virology | A Tradeoff in Replication in Mosquito versus Mammalian Systems Conferred by a Point Mutation in the NS4B Protein of Dengue Virus Type 4 | Dengue virus | *Chlorocebus sabaeus* (Vero cells); *Homo sapiens* (HuH-7 cells); *Aedes albopictus* (C6/36 cells) | Fig 2 | Growth rate; Webplot digitizer |
| Aytay and | 1991 | Journal of | Single Amino Acid | Influenza A | *Bos taurus* (MDBK cells); | Table 2 | Virus titer |

| | | | | | | | |
|---|---|---|---|---|---|---|---|
| Schulze | | Virology | Substitutions in the Hemagglutinin Can Alter the Host Range and Receptor Binding Properties of H1 Strains of Influenza A virus | virus (H1N1) | *Gallus gallus* (CEF cells) | | |
| Scheibner et al. | 2023 | PLOS Pathogens | Phenotypic Effects of Mutations Observed in the Neuraminidase of Human Origin H5N1 Influenza A Viruses | Influenza A virus (H5N1) | *Gallus gallus* (CEK cells); *Homo sapiens* (NHBE cells) | Fig. 1 | Growth rate; Webplot digitizer |
| Arias Goeta et al. | 2014 | Infections, Genetics and Evolution | Chikungunya Virus Adaptation to a Mosquito Vector Correlates with only Few Point Mutations in the Viral Envelope Glycoprotein | Chikunguny a virus | *Aedes aegypti; Aedes albopictus* | Fig.2, Fig. 5 and Fig. 6 | Virus titer growth rate |
| Vasilakis et al. | 2009 | PLOS Pathogens | Mosquitoes Put the Brake on Arbovirus Evolution: Experimental Evolution Reveals Slower Mutation Accumulation in Mosquito than Vertebrate Cells | Dengue virus | *Homo sapiens* (HuH-7 cells); *Aedes albopictus* (C6/36 cells) | Fig. 3 and Fig.5 | Growth rate; webplot digitizer |
| Fitzsimmo ns et al. | 2018 | PLOS Biology | A Speed Fidelity Trade Off Determines the Mutation Rate and Virulence of an | Poliovirus | *Homo sapiens* (Hela cells); *Mus musculus* (3T3 cells) | Fig.1 | Growth rate; webplot digitizer |

| | | | RNA Virus | | | | |
|---|---|---|---|---|---|---|---|
| Coffey and Vignuzzi | 2011 | Journal of Virology | Host Alternation of Chikungunya Virus Increases Fitness while Restricting Population Diversity and Adaptability to Novel Selective Pressures | Chikunguny a virus | *Homo sapiens* (Hela cells); *Aedes albopictus* (C6/36 cells) | Fig.1 | Fitness assays ; webplot digitizer |
| Greene et al. | 2005 | Journal of Virology | Effect of Alternating Passage on Adaptation of Sindbis Virus to Vertebrate and Invertebrate Cells | Sindbis virus | *Mesocricetus auratus* (BHK cells); *Aedes albopictus* (C6/36 cells) | Fig.3 and Fig.4 | Fitness assays; webplot digitizer |
| Furio et al. | 2012 | Journal of Virology | Relationship between within Host Fitness and Virulence in the Vesicular Stomatitis Virus: Correlation with Partial Decoupling | Vesicular stomatitis virus | *Mesocricetus auratus* (BHK21 cells); *Mus musculus* | Table 3 and Table 2 | Fitness assays |
| Van Slyke et al. | 2012 | Virology | Point Mutations in the West Nile Virus (Flaviviridae; Flavivirus) RNA Dependent RNA Polymerase Alter Viral Fitness in a Host-Dependent Manner in Vitro and in Vivo | West Nile virus | *Chlorocebus sabaeus* (Vero cells); *Aedes albopictus* (C6/36 cells); *Culex tarsalis* (CxT cells); *Gallus gallus* (DF-1 cells); *Gallus gallus* | Fig. 3 and Fig.5 | Growth rates; Webplot digitizer |
| Weaver et al. | 1999 | Journal of Virology | Genetic and Fitness Changes Accompanying Adaptation of an Arbovirus to Vertebrate and Invertebrate Cells | Eastern equine encephalitis virus | *Mesocricetus auratus* (BHK cells); *Aedes albopictus* (C6/36 cells); *Chlorocebus sabaeus* (Vero cells); *Anopheles albimanus* | Fig.1 and Fig. 2 | Virus titer fitness assays; Webplot digitizer |

| | | | | | | | |
|---|---|---|---|---|---|---|---|
| | | | | | (pericardial cells) | | |
| Hashemi et al. | 2018 | Journal of Virology | Mutations in the Non Structural Protein Coding Sequence of Proto Parvovirus H1PV Enhance the Fitness of the Virus and Show Key Benefits Regarding the Transduction Efficiency of Derived Vectors | H1 parvovirus (H1PV) | *Homo sapiens* (NB324K cells); *Homo sapiens* (Hela cells); *Rattus norvegicus* (RG2 cells); | Fig.2 | Growth curve; webplot digitizer |
| Tsetsarkin et al. | 2007 | PLOS Pathogens | A Single Mutation in Chikungunya Virus affects Vector Specificity and Epidemic Potential | Chikunguny a virus | *Mesocricetus auratus* (BHK21 cells); *Aedes albopictus* (C6/36 cells); *Aedes aegypti; Aedes albopictus* | Fig. S3, Table S2, Table S1, Fig. 6, Fig. S2 | Growth rate; webplot digitizer |
| Obadan et al. | 2019 | Journal of Virology | Flexibility In Vitro of Amino Acid 226 in the Receptor-Binding Site of an H9 Subtype Influenza A Virus and Its Effect In Vivo on Virus Replication, Tropism, and Transmission | Influenza A virus (H9N1 and H9N2) | *Canis familiaris* (MDCK cells); *Gallus gallus* (DF-1 Cells ) | Fig. 3 | Growth curve; webplot digitizer |
| Lemos et al. | 2020 | Journal of Virology | Two Sides of a Coin: a Zika Virus Mutation Selected in Pregnant Rhesus Macaques Promotes Fetal Infection in Mice but at a Cost of Reduced Fitness in Nonpregnant Macaques | Zika virus | *Aedes albopictus* (C6/36 cells); *Chlorocebus sabaeus* (Vero cells) | Fig. 3 | Growth curve; webplot digitizer |

| | | | | | | | |
|---|---|---|---|---|---|---|---|
| | | | and Diminished Transmissibility by Vectors | | | | |
| Hillung et al. | 2015 | Phil Trans B | Evaluating the within Host Fitness Effects of Mutations Fixed during Virus Adaptation to Different Ecotypes of a New Host | Tobacco etch virus | *Arabidopsis thaliana* (Di-2); *Arabidopsis thaliana* (Wt-1); *Arabidopsis thaliana* (Ei-2); *Arabidopsis thaliana* (Ler-0); *Arabidopsis thaliana* (St-0) | Table 3 | Fitness assays |
| Moreno Perez et al. | 2016 | Journal of Virology | Mutations That Determine Resistance Breaking in a Plant RNA Virus Have Pleiotropic Effects on Its Fitness That Depend on the Host Environment and on the Type, Single or Mixed, of Infection | Pepper mild mottle virus | *Capsicum annuum* cv. Dulce Italiano (L+/L+); *Capsicum annuum* cv. Yolo Wonder (L1/L1); *Capsicum frutescens* cv. Tabasco (L2/L2); | Table 2 | Fitness assays |
| Lalic et al. | 2011 | PLOS Pathogens | Effect of Host Species on the Distribution of Mutational Fitness Effects for an RNA Virus | Tobacco etch virus | *Nicotiana tabacum; Nicotiana benthamiana; Datura stramonium; Capsicum annuum; Solanum lycopersicum; Helianthus annuus; Gomphrena globosa; Spinacea oleracea* | Fig. 3 | Fitness assays |
| Bera et al. | 2018 | Journal of Virology | Analysis of Fitness TradeOffs in the Host | Tobacco mild green | *Capsicum annuus* cv. Doux des Landes; | Table 6 | Fitness assays |

| | | | | | | | |
|---|---|---|---|---|---|---|---|
| | | | Range Expansion of an RNA Virus, Tobacco Mild Green Mosaic Virus | mosaic virus | *Nicotiana glauca* | | |
| Moury et al. | 2011 | Molecular Biology and Evolution | dN/dS Based Methods Detect Positive Selection Linked to Tradeoffs between Different Fitness Traits in the Coat Protein of Potato virus Y | Potato virus Y | *Nicotiana tabacum* cv. Xanthi; *Solanum tuberosum* cv. Bintje | Fig. 2A | Growth rate; webplot digitizer |
| Cervera et al., | 2016 | Journal of Virology | Effects of Host Species on Topography of the Fitness Landscape for a Plant RNA Virus | Tobacco etch virus | *Arabidopsis thaliana; Nicotiana tabacum* | Fig.2 | Fitness assays; webplot digitizer |
| Ayme et al. | 2006 | Molecular Plant Microbe Interactions | Different Mutations in the GenomeLinked Protein VPg of Potato virus Y Confer Virulence on the pvr23 resistance in pepper | Potato virus Y | *Nicotiana clevelandii; Capsicum spp.* cv. Yolo Wonder | Table 4 | Fitness assays |
| Martinez Turino et al. | 2021 | Microorgani-sms | Virus Host Jumping Can be Boosted by Adaptation to a Bridge Plant Species | Plum pox virus | *Arabidopsis thaliana; Nicotiana clevelandii* | Fig. 1 and S3 | Virus accumulation; Image J |

**Table S2**. Adherence to PRISMA-EcoEvo (version 1.0) guidelines for reporting practices in meta-analyses (O'Dea et al., 2021).

| Item No. | Item name | Met criteria? | Explanation |
|---|---|---|---|
| 1 | Title and abstract | Yes | We mention "meta-analysis" both in the title and abstract. |
| 2 | Aims and questions | Yes | We mention our research goals in the introduction and abstract. |
| 3 | Review registration | No | We did not register our study's aims or hypotheses before starting our analysis. |
| 4 | Eligibility criteria | Yes | We include our literature search criteria in the methodology. |
| 5 | Finding studies | Yes | We list all articles our data was sourced from in the supplementary file Table S1. Additionally, we provide our filtering keywords in the methodology. |
| 6 | Study selection | Yes | We read all articles our literature initially identified and decided whether to include based on our criteria. Data collection was initially performed by Muller and McDowell. Wang joined and double checked the data and added more datasets. Rasmussen supervised data collection and double checked data. |
| 7 | Data collection process | Yes | Where in each study data was extracted from is described in supplementary Table S1. Data collection was performed at the same time when the article was initially reviewed. |
| 8 | Data items | Yes | We only sourced the data from published articles, either tables, figures, or supplementary files. We provide additional information on how the data was extracted and analyzed in supplementary Table S1. |
| 9 | Assessment of individual study quality | Yes | All studies were reviewed to see if fitness effects were measured using appropriate standardized methods. |
| 10 | Effect size measures | Yes | Our effect sizes correspond to the reported fitness effect of each mutation in each hose. How fitness effects were standardized and compared is described in the Methods section. |

| | | | |
|---|---|---|---|
| 11 | Missing data | Yes | If mutational fitness effects were missing from a particular host, that host was excluded from our analysis. |
| 12 | Meta-analytic model description | Yes | The linear and mixed linear regression models we used to analyze how fitness effects vary by mutational, viral and host features while accounting for differences between studies (random effects) are described in the Methods. |
| 13 | Software | Yes | All software and packages used to analyze the data are described in the Methods. |
| 14 | Non-independence | Yes | Some of the mutational fitness effects may be non-independent e.g. multiple fitness effects from the same study. Likewise fitness effects may be expected to (co-)vary based on the phylogenetic relationships between viruses and hosts. We account for these sources of non-independence by grouping fitness effects by study, the taxonomic family of viruses, and the phylogenetic relationships (i.e. divergence times) among hosts. |
| 15 | Meta-regression and model selection | Yes | The moderators or features we included in our regression analysis were chosen based on their importance to viral biology or those known from the literature to impact the distribution of fitness effects. While no formal model selection process was performed, we typically fit several alternative models with and without different features to ensure our overall results were robust to correlations between features. |
| 16 | Publication bias and sensitivity analyses | Yes | There is undoubtedly a publication bias resulting in certain viruses or viral families being overrepresented in our analysis. We accounted for these biases by ensuring our main results about the distribution of pleiotropic effects hold more generally and not just for overrepresented taxa. We also performed a sensitivity analysis to ensure our main results about the distribution of pleiotropic effects are robust against varying levels of measurement error in reported fitness effects. |
| 17 | Clarification of post hoc analyses | Yes | We explain when certain analyses (e.g. the inclusion of phylogenetic divergence times between hosts) were performed post hoc based on |

| | | | |
|---|---|---|---|
| | | | prior analysis of the same data. |
| 18 | Metadata, data and code | Yes | The extracted fitness effects and associated metadata are all available at: https://github.com/davidrasm/PleiotropyMetaAnalysis |
| 19 | Results of study selection process | Yes | We report the number of studies included/excluded. |
| 20 | Sample sizes and study characteristics | Yes | The total number of studies, viruses, hosts and mutations are reported in the Results. The number of mutations belonging to each group or subset of viral, host and mutational types are reported in Tables 1, 2, 4 and 5. |
| 21 | Meta-analysis | Yes | We report mean fitness/pleiotropic effects as well as regression coefficients quantifying the impact of different features on fitness effects and their associated confidence intervals. |
| 22 | Heterogeneity | Yes | The variability in fitness effects due to between-study variation was computed using intraclass correlation coefficients (ICC) values. |
| 23 | Meta-regression | Yes | We report regression coefficients and confidence intervals for the effect of features of mutational fitness effects and the magnitude/sign of pleiotropy. These statistics are also reported for interaction effects in our linear models |
| 24 | Outcomes of publication bias and sensitivity analyses | Yes | We performed a sensitivity analysis to see how robust our main results are to different levels of simulated measurement error in the reported fitness effects (see Methods). |
| 25 | Discussion | Yes | We summarize our main findings and compare these results with previous work. We also discuss limitations including potential publication biases. |
| 26 | Contributions and funding | Yes | Author contributions and funding information is provided. |
| 27 | References | Yes | We provide a full list of references to all included studies. |

**Table S3:** Mutational fitness effects in single hosts grouped by viral family. Regression coefficients, P-values and 95% CIs are estimated under a mixed effect linear model that included a random effect (intercept) for each study.

| Variable | N | Group mean | Estimated coefficient | P-value | 95% CI |
|---|---|---|---|---|---|
| Intercept | 457 | - | -0.688 | 0.65 | -1.838 : 0.463 |
| Flaviviridae | 34 | -0.1 | 0.589 | 0.386 | -0.743 : 1.920 |
| Microviridae | 103 | -1.09 | -0.425 | 0.553 | -1.830 : 0.980 |
| Orthomyxoviridae | 44 | 0.35 | 0.624 | 0.392 | -0.805 : 2.054 |
| Parvoviridae | 9 | 0.0001 | 0.688 | 0.432 | -1.026 : 2.401 |
| Picornaviridae | 2 | 0.0048 | 0.692 | 0.512 | -1.379 : 2.764 |
| Potyviridae | 187 | 0.16 | 0.916 | 0.155 | -0.345 : 2.177 |
| Rhaboviridae | 2 | -1.59 | -0.903 | 0.393 | -2.975 : 1.169 |
| Togaviridae | 28 | 0.24 | 1.184 | 0.078 | -0.132 : 2.501 |
| Virgaviridae | 12 | -0.23 | 0.33 | 0.669 | -1.183 : 1.843 |

**Table S4:** Sensitivity of the distribution of pleiotropic effects to measurement error in reported fitness effects. Different levels of error were simulated by adding error terms sampled from a Normal distribution with varying standard deviation σ to the observed fitness effects. Fraction +/- refers to all mutations with antagonistic effects between hosts whereas fraction +/+ refers to the fraction of mutations with unconditionally beneficial effects. The $\chi^2$ statistic corresponds to the test for the independence of the sign of fitness effects between hosts. Reported statistics represent the mean of 1,000 simulations performed at each level of error (σ).

| Error std dev | Corr coeff | Fraction +/- | Fraction +/+ | $\chi^2$ | $\chi^2$ p-value |
|---|---|---|---|---|---|
| 0 | 0.32 | 0.43 | 0.14 | 7.19 | 0.126 |
| 0.01 | 0.32 | 0.41 | 0.13 | 8.59 | 0.085 |
| 0.05 | 0.32 | 0.42 | 0.14 | 8.85 | 0.095 |
| 0.1 | 0.32 | 0.43 | 0.16 | 9.24 | 0.102 |
| 0.25 | 0.31 | 0.44 | 0.19 | 9.33 | 0.124 |
| 0.5 | 0.27 | 0.45 | 0.21 | 8.38 | 0.172 |
| 1 | 0.18 | 0.47 | 0.23 | 6.13 | 0.296 |
